# Warming-induced excess deaths of infected animals depend on pathogen kingdom and evolutionary history

**DOI:** 10.1101/2024.04.16.589701

**Authors:** Jingdi Li, Nele Guttmann, Georgia Drew, Tobias Hector, Justyna Wolinska, Kayla King

## Abstract

Climate change is causing extreme heating events. Simultaneously, climate change and human activities are leading to more prolonged and intense infectious disease outbreaks. The extent to which warming and infection may together impact host species persistence is, however, unclear. Using a meta-analysis of >190 effect sizes representing 101 ectothermic animal host-pathogen systems, we provide broad evidence that experimentally increased temperatures drove higher pathogen virulence, specifically pathogen-induced host mortality. Such pattern was mainly driven by excess host death caused by bacterial infections combined with warming, particularly if the pathogenic bacteria were naturally established within the host species, though novel infections without known host-pathogen evolutionary history were more lethal at lower temperatures. Importantly, larger temperature increases were associated with more host deaths hinting at the escalating threat for animal species as the world continues to warm. We found that the virulence of fungal pathogens increased only when temperatures were shifted upwards towards their thermal optimum. The magnitude of these effects was not impacted by host life-stage, immune complexity, or variable experimental protocols. Overall, our findings reveal distinct patterns of pathogen virulence change under warmer temperatures, suggesting that the impact of global warming on infectious disease outcomes would depend on pathogen traits (taxonomic kingdom, thermal tolerance) and host-pathogen evolutionary history.

**Author Summary:** Human-induced climate warming is one of the biggest challenges in our times. Simultaneously, climate change is associated with more intense infectious disease outbreaks, suggesting that temperature rises also influence disease dynamics. Growing numbers of studies have investigated the effect of warming on disease severity (or pathogen virulence) in different animal host-pathogen systems. However, individual studies did not always agree with each other, and how increased temperature and pathogen infection together impact animal survival remains unclear. Here, we resolved this uncertainty by conducting a meta-analysis of >190 effect sizes representing 101 animal host-pathogen systems. We provided broad evidence that, higher temperatures caused more deaths of infected animals, particularly for animals with bacterial infections under warmer conditions. We found that larger temperature rises were associated with more animal deaths, suggesting the increased threat for host species as the world continues to warm. We also found that pathogenic fungi were more sensitive to heat than bacterial pathogens, and temperature changes the virulence of fungal pathogens in relation to their thermal optimum.

## Introduction

Climate change is resulting in more extreme heating events [1]. Temperature affects all aspects of biology, directly or indirectly impacting the physiology and life-history of all organisms [2]. Typically for ectotherms, environmental warming changes their body temperatures, influencing their physiological sensitivity and fitness [3]. Shifting global temperature patterns are also changing the geographic distribution of infectious diseases, generating novel transmission opportunities [4,5].

Positive relationships between warmer temperatures and disease severity (or pathogen virulence) have been detected in a diversity of systems [6–11]. Rising temperature can alter animal immune responses, leading to increased susceptibility to infections [12–15]. Moreover, by causing stress and disrupting homeostasis, pathogen infection can reduce the upper temperatures tolerated by hosts, contributing to worsened health outcomes under warming [16]. Temperature also directly mediates pathogen virulence [17–21]. Higher virulent temperatures can upregulate the expression of virulence factors in the coral pathogen *Vibrio coralliilyticus* [20]. For human pathogen *Shigella sonnei*, higher temperature enhances its immune evasion abilities by increasing virulence protein synthesis [21]. Warming may also enhance pathogen metabolism, resulting in faster within-host growth and greater host harm. This might be especially true for many bacterial pathogens, as their population growth rates can increase exponentially with temperature, up to their thermal limit [22]. Yet in other host-pathogen systems virulence is unaffected or even reduced by warming, emphasizing complex interactions between temperature, biology and disease outcomes [23–28].

Both host and pathogen responses to temperature changes depend on their thermal tolerance, which is characterized by the critical thermal maximum, minimum and optimum - the upper, lower and optimal temperatures at which the organism can function [3,29,30]. A mismatch in host and pathogen thermal optima can change infection outcomes, such that cold-adapted pathogens might be less harmful in a warm-adapted hosts at higher temperatures [31,32]. Also, eukaryotes normally have lower thermal tolerance compared to prokaryotes [33]. This difference might render eukaryotic pathogens (e.g. fungi, nematodes) more sensitive to warmer temperatures. Clearly not all host-pathogen interactions will be affected equally by climate change, and it is unclear which outcome will apply to which type of infection [23,24].

Understanding the broad ecological consequences of elevated temperature on pathogen virulence (or pathogen-induced host death herein), is crucial for projections of future wildlife disease dynamics and species persistence. If elevated temperature generally exacerbates the severity of infection, climate change could worsen disease outcomes for the wildlife. Projections of species persistence in a changing world might need to account for rates of infection and disease severity [8,34]. However, if virulence is broadly unaffected or context-dependent in specific types of infection [23], the health risks posed by warming and infectious disease could be considered on a case-by-case basis. Any broad relationship between warming and virulence would have significant consequences for animal conservation [34,35] and potentially human health [4,36].

In this study, we conducted a meta-analysis to investigate the generality of the relationship between elevated temperature and pathogen virulence. Are warmer hosts also sicker? We searched the published literature for experimental studies and included a breadth of ectothermic animal host-pathogen systems. We used the most widely reported quantitative form of virulence: pathogen-induced host death [37]. We focused on host death under infection and warming as this measure was relevant for consideration in projections of species extinction and biodiversity loss [38]. We calculated effect sizes via relative risk ratios of death at experimentally high and low temperatures. We separately collected thermal optima data for all the pathogens included. We tested whether biological traits (e.g. host type, pathogen type, host-pathogen evolutionary history, pathogen thermal optimum), the scale of temperature increase and experimental variables (pathogen inoculation method, pathogen dosage, infection exposure/measurement time) could explain variation in the temperature-virulence relationship.

## Material and methods

### Literature search and data collection

A first-round literature search was performed on March 2021 on Web of Science Core Collection using keywords ‘parasit*’ OR ‘pathogen*’, paired with ‘temperature’ AND ‘virulen*’, to identify studies reporting the effect of temperature on pathogen virulence. Then a second-round search using the same keywords was conducted on April 2022 to update the dataset. We used host mortality to quantify pathogen virulence. We excluded studies on parasitoids as host mortality under infection would always be 100%. Our main inclusion criteria were: (i) the pathogen was tested in living animal hosts rather than on cells or dead host tissues; (ii) more than one temperature was used in host infection experiment and temperature was the only changing variable in the tested comparisons; (iii) pathogen virulence was measured by assessing host mortality rate, under different temperatures; (iv) host mortality data (mean mortality rate and sample size, sample size is defined as the number of individual host used in each infection treatment) were available publicly or shared by the author by May 2022.

Following our inclusion criteria, we retrieved data from a total of 60 research articles (PRISMA flow-chart see Fig. S1, with PRISMA-EcoEvo Checklist included), published between 1996 and 2021 (summary of studies and moderator variables are provided in Table S1). To maximise the ecological relevance, we imposed a series of restrictions to exclude data from extreme high or low temperatures and extreme high pathogen dosages unlikely to happen in nature. Firstly, if more than five temperatures were tested, data were extracted from the second lowest and second highest temperatures (when between two to four temperatures were tested, data were extracted from the highest and lowest temperatures). Secondly, if multiple pathogen dosages were used, data of the middle value (or middle minus one value, in case of even number) was extracted. Thirdly, host mortality data was extracted from the end time point of the assay. However, if mortality began to appear in controls, data were extracted from the timepoint with control mortality close to or at zero. As a result, we focused on infection-induced host mortality, rather than death due to temperature alone.

We ensured that all host mortality in the pathogen treatment groups was due to infection rather than other reasons. Hosts not exposed to pathogens had no mortality (e.g. Jiang et al. 2013), very low mortality (<10%, e.g. Brand et al. 2016) or host mortality in the infection treatment was corrected for control mortality following Abbott’s (1952) formula (e.g. Bugeme et al. 2008). Data on host mortality in individual articles were extracted manually from text and tables in the main text or supplementary information, figures by using WebPlotDigitizer [39], or raw data made available from the authors by request. Fifteen effect sizes from six studies were discarded due to large host mortality in control groups.

When host mortality was expressed as a percentage, we estimated the number of alive and dead individuals in both temperature groups by multiplying percentage of host mortality by the number of host individuals tested. We otherwise obtained the number of alive and dead hosts from the raw data sent by authors. These data were used to calculate the risk ratio (RR) effect size, using *escalc* function in ‘metafor’ R package [40]. RRs represent the relative risk of warming on infected hosts’ survival. An overall total of 192 effect sizes from 60 studies were used across 56 pathogens and 50 host species (101 host-pathogen combinations, Fig. S2).

### Moderator variables

Host, pathogen and experimental variables were included to assess their impact on pathogen virulence when temperature rose. To explore whether pathogen type could predict effect size estimates, we recorded whether the pathogen used in the study was bacteria, fungi, nematode, virus, or protist. As the number of effect sizes in protist was low (1 ES for protist), our analysis focused on bacteria, fungi, nematode and virus pathogens.

We collected data on the optimal growth temperature (T_opt_) for well-studied pathogens as virulence can correlate with pathogen growth rates [41]. These data were collected for 39 pathogen species, included in the meta-analysis, from published literature (Table S3, T_opt_ for *in vitro* growth/fitness were measured for non-virus pathogens). Using known thermal optimums for these pathogens, we were able to define whether experimental warming was towards or away from their T_opt_. “Towards T_opt_” was used when the temperature range in the study was below T_opt_. “Away from T_opt_” was used when the study temperature range was above T_opt_. When the low and high temperatures in the study straddled T_opt_, we used “Range includes T_opt_”.

To test whether the response to increased temperature depended on host-pathogen evolutionary history, we defined a moderator based on whether the interaction was “established” (system originally collected from the wild and with previous evolutionary history (see Bally and Garrabou 2007 as an example), “semi-established” (host infected by the pathogen in the wild, but combination in the study were not co-isolated (see Wekesa et al. 2010 as an example), and “novel” (no known record of an infection between the pathogen and the host in nature, see Ekesi et al. 1999 as an example). The “novel” associations might represent opportunistic infections or a host shift for the pathogen.

Host traits including host life-stage (“adult”, “larva”, “pupae” and “juvenile”), host source (“collected from wild”, “lab-reared”) and host type with different immune complexity (“vertebrate”, “invertebrate”) were also tested as moderator variables. Lastly, we collected and evaluated experimental variables including temperature span (difference between high temperature and low temperature used in the study), infection exposure duration (measured by number of days), pathogen dosage (within each pathogen type) and pathogen inoculation method (“injection”, “not injection”). These experimental protocols were evaluated given their potential for a bias in extrapolations to nature [42].

### Statistical analysis

By fitting sample size or correction method as a moderator, we checked that neither sample size nor the methods used to correct for control mortality influenced the effect sizes (sample size p=0.9630, correction method p>0.52 for all levels). All categorical moderators were assessed for multicollinearity by calculating the variance inflation factor (VIF, *VIF* function in ‘regclass’ R package) when fitted together in a regression model and no multicollinearity was observed (Table S4, GVIF^(1/(2*Df)) <= 2.5 for all moderators included). Correlation of continuous variables were assessed for Pearson correlation (cor.test function in R) and the possibility of multicollinearity was excluded (Table S4, p>0.07 for all pairwise correlations).

We coded the data, whereby RR>1 (logRR>0) indicated greater host mortality at higher temperatures (higher risk associated with higher temperatures) and RR<1 (logRR<0) indicated lower host mortality at higher temperatures. A relationship in which temperature has no effect on pathogen virulence was indicated by RR=1 (logRR=0). We fitted a multi-level random-effects model using *rma.mv* function in ‘metafor’ R package to estimate mean effect sizes across all the studies. We used host species, pathogen species and study as random factors to account for statistical non-independence due to repeated measures within species and studies. We did not include the phylogenetic tree of either host or pathogen species as a random factor. For many cases, we could not resolve species-level phylogeny from published literature, and replacing those with their relatives would introduce more bias.

For categorical moderators, we first conducted sub-group analysis to calculate summary effect size for each sub-group. We then assessed whether there was significant difference between different sub-group levels by fitting the moderator in multilevel random effects meta-regression models (*rma.mv* function in R). For continuous moderators, we fitted the moderator in multilevel random effects meta-regression models (*rma.mv* function in R) with the effect sizes as response variables.

Separate tests of interactions between moderators were driven by *a priori* hypotheses (all hypotheses and model results in Table S4).

## Results and Discussion

### More death of infected hosts at hotter temperatures

We firstly tested whether hotter temperatures generally increase host death (i.e. the proportion of dead individuals in the infected host population). We incorporated data from 60 studies which conducted infection experiments at different temperatures. These studies included 50 ectothermic animal host species and 56 pathogen species (resulting in 101 analysed host-pathogen combinations, see Fig. S2), from which we calculated 192 effect sizes (ESs). Our data set contained a variety of pathogen types including bacteria, fungi, nematodes, and viruses from diverse geographic regions (Fig. S3). We included diverse host-pathogen systems spanning terrestrial (42 ESs) and aquatic habitats (150 ESs). Terrestrial hosts mostly consisted of Insecta (125 ESs), and aquatic mostly consisted of fish (17 ESs) and mollusca (14 ESs) (Fig. S2). Overall, bacterial pathogens were mostly studied in aquatic animals, while most fungal pathogens as well as all the nematoda pathogens were tested in insecta hosts (Fig. S2).

Under warming conditions, the rate of uninfected host mortality across studies remained zero or low (<10%). In contrast, infected hosts suffered greater death rates with increase in temperature (Fig. 1A, logRR>0, RR=1.47, p=0.0046). Previous findings that infection can reduce the ability of hosts to tolerate elevated temperatures [16,43,44] may reflect this increase in pathogen virulence.

**Figure 1.**
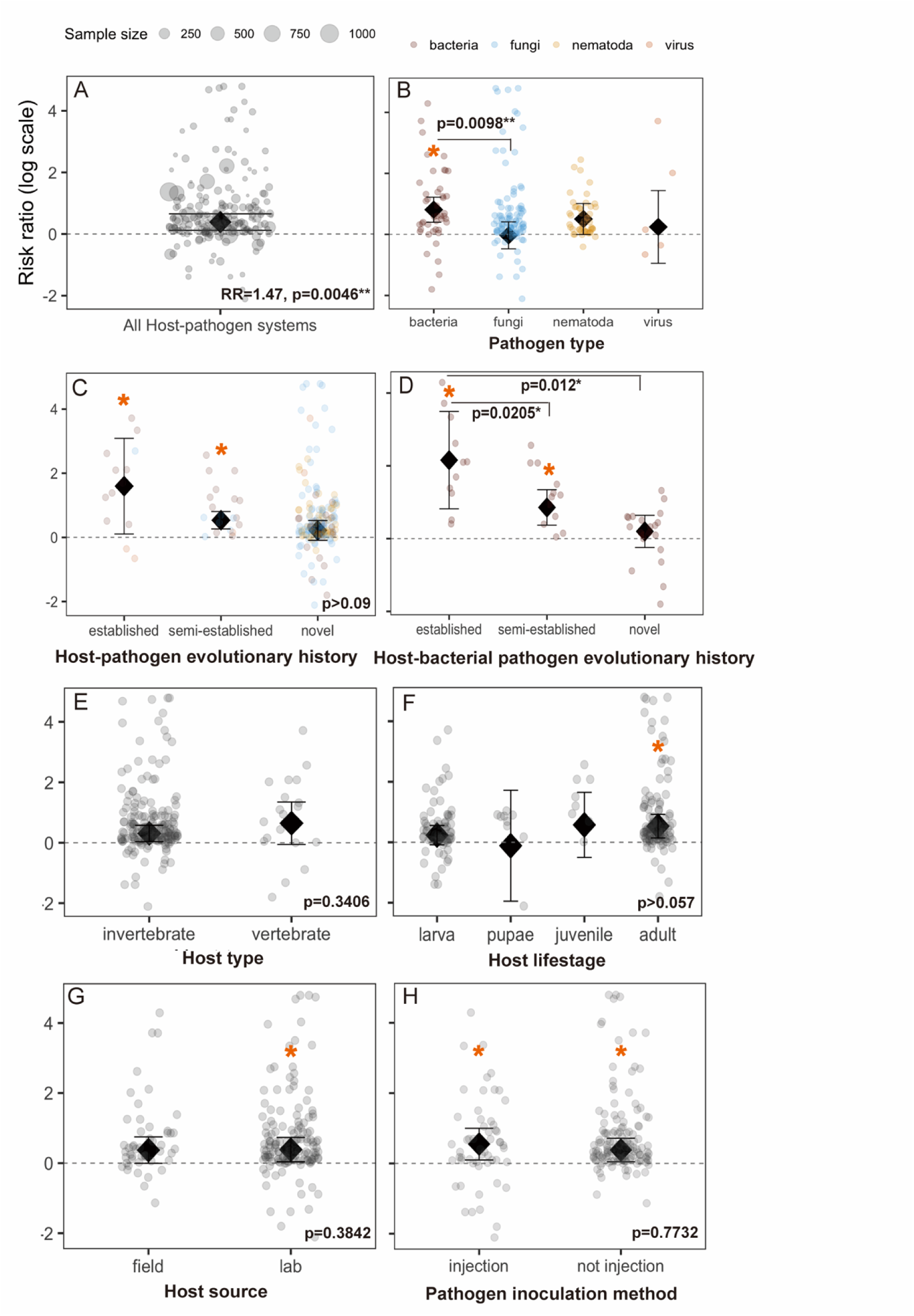
Results of the meta-analysis on the impact of heating on pathogen-induced host death. (A) Summary effect size (Risk ratio, RR) from overall meta-analysis is significantly >1 (logRR>0, indicating positive relationship between temperature and virulence). Bold black diamond with 95% confidence interval represents summary effect size, and individual effect size are displayed as jittered points. Point size indicates sample size (number of host individuals in the infection treatment) used for effect size calculation. The summary RR with the corresponding p-value is shown in the bottom right corner of the plot. The presence of the effect differed by (B) pathogen type and (D) host-pathogen evolutionary history for bacterial infections, but less pronounced when combined all effect sizes across pathogen types (C). The presence of an effect was not impacted by (E) host type with varying immune complexity, (F) host early or adult life-stage, (G) host source being wild-collected or lab-reared or (H) pathogen was inoculated by injection or other means. Orange star is shown above the subgroups that have summary RR significantly >1. P-values shown on the bottom right corner indicate significance level of subgroup comparisons. In (B-D), jittered points are colored by pathogen type.

We subsequently investigated whether biological factors could explain the variation in effect sizes, using sub-group analysis and meta-regressions. We specifically tested whether the relationship between temperature and virulence differed between (i) pathogen types (bacteria, fungi, nematoda, viruses), (ii) host-pathogen evolutionary history (established vs. novel), (iii) host types and immune complexity (vertebrate vs. invertebrate), (iv) host life-stage (e.g. adult vs. larva), and (v) wild-collected vs. lab-reared hosts.

We found that the impact of higher temperatures on host death differed among pathogen types. Bacterial pathogens exhibited the strongest and most positive effect sizes (Fig. 1B, bacteria RR=2.25, p=0.0001; fungi RR=0.97, p=0.8930, nematoda RR=1.66, p=0.0523, virus RR=1.28, p=0.6868; bacteria vs. fungi p=0.0098, p>0.09 for all other comparisons). As most of the effect sizes for bacteria were calculated from aquatic hosts, our results suggest that bacterial pathogens might pose significant greater risk under warming, to aquatic hosts such as fishes. Increased disease outbreaks for aquatic animals, especially fishes, has been an overwhelming issue for the aquaculture industry, causing considerable economic damage and threats to human health [45]. Bacterial diseases are common in aquaculture, we suggest that future disease management should also consider the aggravating effects of warming on the severity of epidemics.

Our results showed that host death increased with warming in established, but not novel host-pathogen systems (Fig. 1C, established RR=4.94, p=0.0357; semi-established RR=1.71, p=0.0001, novel RR=1.2, p=0.1639). However, no significant difference was found between “established” and “novel” subgroups (Fig. 1C, p>0.09). Since the interactive effect was detected between pathogen type and evolutionary history (p=0.0014 for “novel”:”virus”, Table S4), we then assessed the effect of host-pathogen evolutionary history within each pathogen type. We found that the effects of excess host death in established, but not novel systems were the most pronounced in host-bacterial pathogen systems (Fig. 1D, established RR=8.64, p=0.0016; semi-established RR=2.35, p=0.0006; novel RR=1.22, p=0.3794; established vs. novel p=0.0120). These effects were significant regardless of host type (p=0.0664 for “host-pathogen system”:“host type”, Table S4). We also found that host mortality in novel systems was already higher than that in established associations, at low/baseline temperature (Fig. S4A, C, p=0.0007). This high level of virulence might be due to pathogen maladaptation when it shifts to a new host [46]. These results suggest that emerging infections might already be highly virulent at present ambient temperatures, and less likely to be worsened by temperature increases specifically during global climate change. On the contrary, the increased severity of endemic infections should be considered when predicting wildlife persistence under warming.

There was no evidence that the magnitude of the temperature-host death association was influenced by other moderators tested – host life-stage, host type, or host source (Fig. 1E-G, p>0.05 for all comparisons). The overall pattern of increased pathogen virulence under warming was also consistent despite variation in experimental protocols. We tested the impact of these experimental moderators (i) pathogen inoculation method (injection vs. not injection), (ii) pathogen dosage, and (iii) exposure or measurement time points. Host death was increased at hotter temperatures consistently across all protocols investigated (Fig. 1H, pathogen inoculation method: injection: RR=1.73, p=0.0172; not injection: RR=1.46, p=0.0258; comparison: p=0.7732; Fig. S5, pathogen dosage: p>0.05 for all pathogen types; infection exposure time: p=0.8642).

### More host death with larger temperature changes

We tested whether the magnitude of the temperature-death relationship would scale with temperature increases. We fit temperature span as a moderator in a meta-regression. Larger temperature changes in experiments were associated with bigger effect sizes (Fig. 2A, p<.0001), regardless of any biological factors (Table S4, p>0.1 for all interactions). We then assessed the robustness of the empirical evidence by re-analysing a single fungal pathogen species - *Beauveria bassiana*, which had the most data across multiple arthropod host species (25 effect sizes). Similarly, higher levels of host death during infection by this fungal species were associated with greater temperature changes (Fig. S6, p=0.0110).

**Figure 2.**
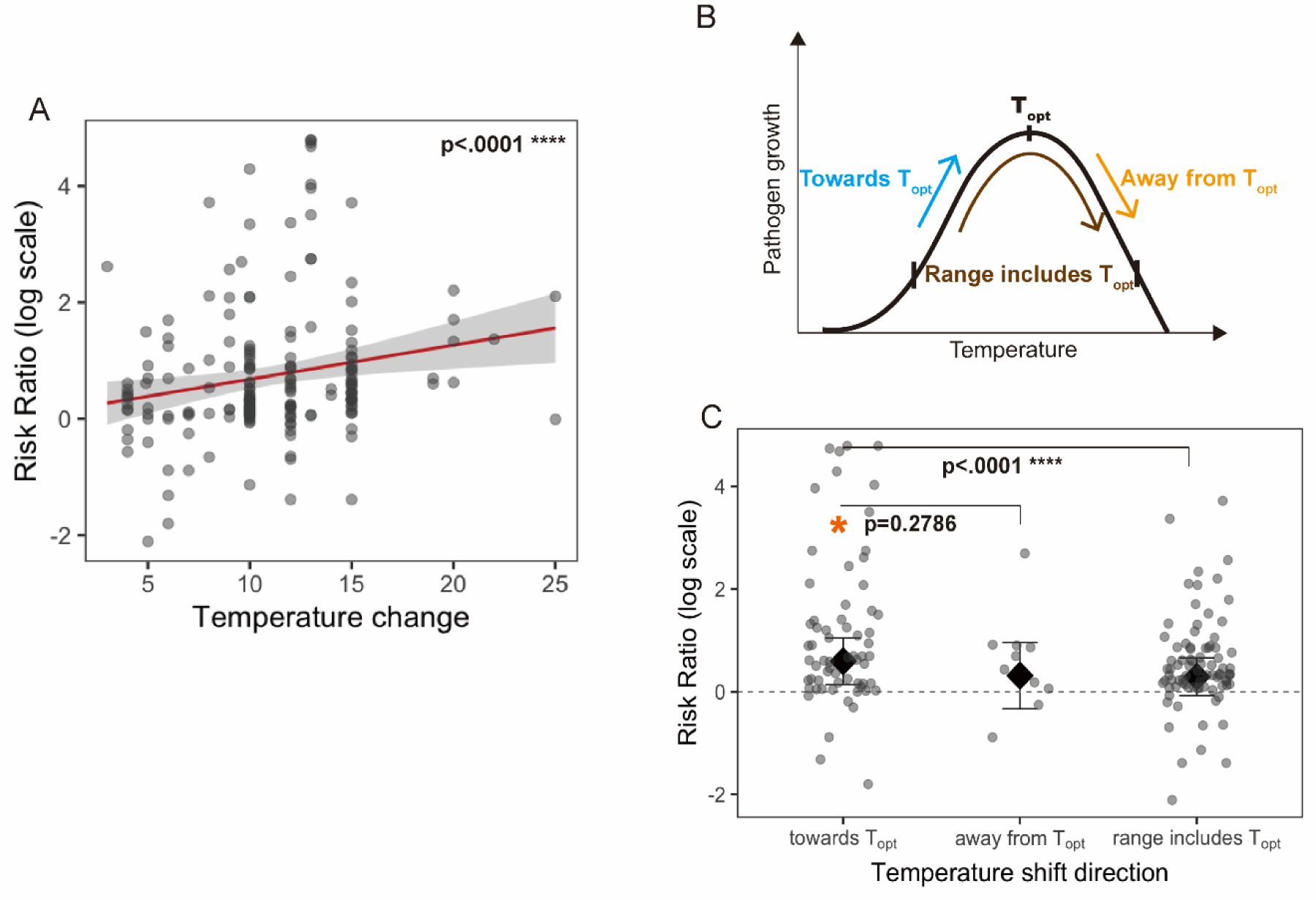
Risk ratio (RR) varied with scale and direction of temperature change. (A) Positive association between scale of temperature change and effect sizes. (B) Schematic illustration of the classification for directions of temperature shift related to pathogen thermal optima (T_opt_). The curve represents a generic thermal performance curve with T_opt_ as the optimal growth temperature. The direction of a temperature shift is represented by arrows. “Towards T_opt_”: the temperature range in the study was below T_opt_. “Away from T_opt_”: the temperature range was above T_opt_. “Range includes T_opt_”: the low and high temperatures in the study straddled T_opt_. (C) Summary RR for subgroups of temperature shift directions. Black diamond with 95% confidence interval represents summary effect size, and individual effect sizes are displayed as jittered points. Asterisk indicates RR significantly > 1 (logRR>0) for subgroup “towards T_opt_”. P-values between subgroups show level of significance.

Experimental temperature increases included in our analysis ranged from 3°C to 25°C. This range is relevant to the upper boundary of the predicted temperature rise on Earth: 4°C increase in annual mean surface temperature by 2100 and 14.1°C increase by 2300 [47]. If no actions are taken to alleviate climate change, our findings suggest that the outcomes of infectious disease could worsen to a greater extent in regions of more intense warming and more frequent extreme thermal events [48]. More studies are needed to investigate whether host responses (i.e., evolutionary adaptation in shorter-lived species, acclimatization in longer-lived species, and migration in both) to warming could help mitigate these negative effects [49].

### Pathogen thermal optima may hint at a mechanism

We hypothesized that the increase in host death may be the result of faster pathogen growth or higher within-host loads at higher temperatures. We estimated pathogen thermal optima (T_opt_) using *in vitro* growth rate data for 39 well-studied species (Table S3). We found that warming increased virulence when temperature was elevated in the direction of pathogen optima, using sub-group analysis on temperature changes relative to T_opt_ (Fig. 2B, Fig. 2C, subgroup “towards T_opt_”: RR=1.81, p=0.0104). This indicated that warmer and more optimal temperatures can promote pathogen growth, leading to heightened pathogen load and greater host harm. The extent of change in host death was significantly different between subgroup “towards T_opt_” and “range includes T_opt_” (Fig. 2C, p<.0001), but not significantly different between subgroup “towards T_opt_” and “away from T_opt_” (Fig. 2C, p=0.2786).

We then compared effect sizes of these subgroups within the same pathogen type. We found that the direction of the temperature shift relative to T_opt_ was more vital for outcomes of fungal infections, compared to other types of pathogens (i.e. bacteria, nematoda). Unlike bacterial infections, fungal pathogens did not result in higher average host death under warmer temperatures (Fig. 1B, RR=0.97, p=0.8930), supporting the observation that fungi are more heat sensitive compared to bacteria [50]. However, host death caused by fungi increased when the temperature was shifted towards fungal T_opt_ (“towards T_opt_”:“fungi” es>0, p=0.0181, Fig. S7). As 92% of fungal effect sizes were calculated from novel host-fungi systems, this result might help predict patterns of emerging fungal diseases. Most fungal species are limited in their capability to grow at elevated temperatures which limits their potential to infect mammals [51]. Our finding suggests that fungal pathogen species could be deadlier if temperatures shifted towards their T_opt_. This is in consistence with previous findings that *Batrachochytrium dendrobatidis* (Bd) – a deadly fungal pathogen driving the extinction of amphibian species – is more prevalent at highland localities where the temperatures are shifted towards the pathogen’s T_opt_ [52]. At higher temperate latitudes, the persistently warmer temperatures caused by climate change are contributing to the ongoing expansion in the ranges of fungal pathogens such as *Coccidioides*, *Blastomyces*, *Histoplasma*, and *Sporothrix* [53,54]. With continued warming in these regions, emerging fungal infections could spread faster and be more severe, posing a greater threat to susceptible and endangered hosts, such as amphibians and reptiles [55].

Apart from the direction of temperature shift, host thermal tolerance and immune response might also play important roles in shaping such infection outcomes. For example, host immunopathology induced by infection can be heightened under warmer conditions, causing host cell and tissue damage and worsening infection outcomes [56,57]. It can be difficult to distinguish the adverse effects of host immunopathology from direct effects of pathogens under warming [58]. Further experiments on the impact of temperature on immune activation and immunopathology are needed. Tackling the impact of global climate change on wildlife health requires an understanding the mechanisms that might increase host mortality during infection when periods of extreme warming hit.

### Conclusions and future perspective

A greater diversity of host-pathogen systems should be included in future experimental tests of the temperature-virulence relationship. While invertebrate hosts included in our analysis ranged from terrestrial (e.g., insects, mite, etc.; 78.1% of the effect sizes) to aquatic (e.g., corals, oysters, etc.; 12% of the effect sizes), there were fewer experimental studies with aquatic vertebrate hosts (fishes, amphibians, 9.9% of the effect sizes) and none with terrestrial vertebrates. Bias towards this limited set of systems could be due to most of our included studies stemming from the field of agriculture (e.g., used Arthropoda hosts) or aquaculture (e.g., used fish hosts). Our study focused on ectotherms in which both the host and pathogens are directly exposed to warming, whether endotherm hosts – with more complex thermal regulation mechanisms – have similar responses requires further study. Indeed, most emerging human pathogens originate in other vertebrates, especially mammals [59–61]. Further research on the impact of higher temperatures in vertebrate-pathogen systems is essential for robust predictions on the outcomes of infections with zoonotic potential under climate change.

Both ecological and evolutionary consequences of elevated temperatures are important for understanding the impacts of warming on ecosystems. Our study revealed temperature’s effect on infection outcomes on an ecological timescale, for which pathogen and warming exposure were not carried across multiple host generations. Over longer evolutionary timescales, the difference between the host’s and pathogen’s ability to acclimate or adapt with thermal changes may be important. Due to their generation times and faster metabolisms, pathogens might adapt more rapidly to temperature shifts than their hosts, leading to increased pathogen growth within hosts [62]. To test whether the pattern on ecological timescales will remain the same across evolutionary timescales, longitudinal studies (perhaps using experimental evolution) could be used to track the potential for host and pathogen adaptation under temperatures relevant to climate change projections. Slow temperature increases may accommodate host acclimation or adaptation thereby mitigate the negative effect of warming on infection outcomes [63].

Parasites and pathogens are ubiquitous. Thus, understanding the impact of warming on the harm caused by infection is critical for projections of animal species persistence under global climate change [64–68]. Our meta-analysis provides evidence that warming, and the degree and direction of temperature increase, can potentially lead to excess mortality in infected animal populations. These patterns indicate distinct consequences of warming on bacterial and fungal infections, and for hosts in different habitats. More intense warming, if concomitant with epidemics caused by bacterial agents, could exacerbate aquatic animal population declines. While emerging fungal diseases for terrestrial animals are more of a concern when temperature rise is more optimal for pathogen growth. There are important consequences in this temperature-virulence relationship for species at risk and ecosystems sensitive to biodiversity loss. Human activities are warming the Earth at an unprecedented rate [69] and leading to an increased frequency and intensity of temperature extremes [47]. The current global effort to restrict warming to 1.5°C above pre-industrial levels by 2050 [70] may also have the unintended benefit of limiting the lethal impacts of infectious diseases.

## Statement of authorship

JL, JW, KCK and GCD designed the study. JL and NG conducted literature review and collected data. JL performed the meta-analysis, with input from JW, KCK, GCD and TEH. JL and KCK wrote the manuscript. All authors contributed to reviewing and editing.

## Data accessibility statement

The data that support the findings of this study are openly available in Figshare at https://doi.org/10.6084/m9.figshare.22060646.v4.

## References

1. Dosio, A., Mentaschi, L., Fischer, E. and Wyser, K., Extreme heat waves under 1.5°C and 2°C global warming, ENVIRONMENTAL RESEARCH LETTERS, ISSN 1748-9326, 13 (5), 2018, p. 054006, JRC108180.

2. Lagerspetz K. Y. (1987). Temperature effects on different organization levels in animals. Symposia of the Society for Experimental Biology, 41, 429–449.

3. Huey R.B. et al. 2012. Predicting organismal vulnerability to climate warming: roles of behaviour, physiology and adaptation. Philosophical Transactions of the Royal Society of London B: Biological Sciences 367:1665–1679.

4. Williams, E. S., Yuill, T., Artois, M., Fischer, J., & Haigh, S. A. (2002). Emerging infectious diseases in wildlife. Revue scientifique et technique (International Office of Epizootics), 21(1), 139–157. 10.20506/rst.21.1.1327

5. Roberts, K. E., Hadfield, J. D., Sharma, M. D., & Longdon, B. 2018. Changes in temperature alter the potential outcomes of virus host shifts. PLoS pathogens, 14(10), e1007185. 10.1371/journal.ppat.1007185

6. Vezzulli, L., Previati, M., Pruzzo, C., Marchese, A., Bourne, D. G., Cerrano, C., & VibrioSea Consortium (2010). Vibrio infections triggering mass mortality events in a warming Mediterranean Sea. Environmental microbiology, 12(7), 2007–2019. 10.1111/j.1462-2920.2010.02209.x

7. Jiang, Q., Shi, L., Ke, C., You, W., & Zhao, J. (2013). Identification and characterization of Vibrio harveyi associated with diseased abalone Haliotis diversicolor. Diseases of aquatic organisms, 103(2), 133–139. 10.3354/dao02572

8. Paull S. H., P. T. Johnson, Experimental warming drives a seasonal shift in the timing of host-parasite dynamics with consequences for disease risk. Ecol. Lett. 17, 445–453 (2014).

9. Mayers TJ, Bramucci AR, Yakimovich KM and Case RJ (2016) A Bacterial Pathogen Displaying Temperature-Enhanced Virulence of the Microalga Emiliania huxleyi. Front. Microbiol. 7:892. doi: 10.3389/fmicb.2016.00892

10. Brand, M. D., Hill, R. D., Brenes, R., Chaney, J. C., Wilkes, R. P., Grayfer, L., Miller, D. L., & Gray, M. J. (2016). Water Temperature Affects Susceptibility to Ranavirus. EcoHealth, 13(2), 350–359. 10.1007/s10393-016-1120-1

11. Bugeme, D. M., Knapp, M., Boga, H. I., Wanjoya, A. K., & Maniania, N. K. (2009). Influence of temperature on virulence of fungal isolates of Metarhizium anisopliae and Beauveria bassiana to the two-spotted spider mite Tetranychus urticae. Mycopathologia, 167(4), 221–227. 10.1007/s11046-008-9164-6

12. Moriyama, M., Ichinohe, T. 2019. High ambient temperature dampens adaptive immune responses in influenza A virus infection. Proceedings of the National Academy of Sciences USA 116, 31183125.

13. Regnier, J.A., and K.W. Kelley. 1981. Heat- and cold-stress suppresses in vivo and in vitro cellular immune responses of chickens. Am. J. Vet. Res. 42:294–299.

14. Lecchi, C., N. Rota, A. Vitali, F. Ceciliani, and N. Lacetera. 2016. In vitro assessment of the effects of temperature on phagocytosis, reactive oxygen species production and apoptosis in bovine polymorphonuclear cells. Vet. Immunol. Immunopathol. 182:89–94.

15. Lacetera, N., U. Bernabucci, D. Scalia, B. Ronchi, G. Kuzminsky, and A. Nardone. 2005. Lymphocyte functions in dairy cows in hot environment. Int. J. Biometeorol. 50:105–110. doi:10.1007/s00484-005-0273-3.

16. Hector, T. E., Hoang, K. L., Li, J., & King, K. C. (2022). Symbiosis and host responses to heating. Trends in Ecology & Evolution, 37, 611– 624.

17. Laine A.L. 2007. Pathogen fitness components and genotypes differ in their sensitivity to nutrient and temperature variation in a wild plant–pathogen association. Journal of Evolutionary Biology. 20:2371–2378.

18. Vale P.F. & T. J. Little. 2009. Measuring parasite fitness under genetic and thermal variation. Heredity 103:102–109.

19. Allen D.E. & T. J. Little. 2011. Dissecting the effect of a heterogeneous environment on the interaction between host and parasite fitness traits. Evol Ecol 25:499–508.

20. Kimes NE, Grim CJ, Johnson WR, Hasan NA, Tall BD, Kothary MH, Kiss H, Munk AC, Tapia R, Green L, Detter C, Bruce DC, Brettin TS, Colwell RR, Morris PJ. 2012. Temperature regulation of virulence factors in the pathogen Vibrio coralliilyticus. ISME J. 6(4):835–46. doi: 10.1038/ismej.2011.154.

21. Lam, O., Wheeler, J., & Tang, C. M. (2014). Thermal control of virulence factors in bacteria: a hot topic. Virulence, 5(8), 852–862. 10.4161/21505594.2014.970949

22. Qiu, Y., Zhou, Y., Chang, Y., Liang, X., Zhang, H., Lin, X., Qing, K., Zhou, X., & Luo, Z. (2022). The Effects of Ventilation, Humidity, and Temperature on Bacterial Growth and Bacterial Genera Distribution. International journal of environmental research and public health, 19(22), 15345. 10.3390/ijerph192215345

23. Lafferty K. D., E. A. Mordecai, The rise and fall of infectious disease in a warmer world. F1000 Res. 5, 2040 (2016).

24. Karvonen A, Rintamaki P, Jokela J, Valtonen ET. Increasing water temperature and disease risks in aquatic systems: Climate change increases the risk of some, but not all, diseases. Int J Parasitol. 2010;40(13):1483–1488. doi: 10.1016/j.ijpara.2010.04.015.

25. Delisle, L., Petton, B., Burguin, J. F., Morga, B., Corporeau, C., & Pernet, F. (2018). Temperature modulate disease susceptibility of the Pacific oyster Crassostrea gigas and virulence of the Ostreid herpesvirus type 1. Fish & shellfish immunology, 80, 71–79. 10.1016/j.fsi.2018.05.056

26. De Silva, P. M., Chong, P., Fernando, D. M., Westmacott, G., & Kumar, A. (2017). Effect of Incubation Temperature on Antibiotic Resistance and Virulence Factors of Acinetobacter baumannii ATCC 17978. Antimicrobial agents and chemotherapy, 62(1), e01514–17. 10.1128/AAC.01514-17

27. Kerchev, I.A., Kryukov, V.Y., Yaroslavtseva, O.N. et al. The first data on fungal pathogens (ascomycota, hypocreales) in the invasive populations of four-eyed fir bark beetle Polygraphus proximus Blandf.. Russ J Biol Invasions 8, 34–40 (2017). 10.1134/S2075111717010040

28. López, J. R., Lorenzo, L., Marcelino-Pozuelo, C., Marin-Arjona, M. C., & Navas, J. I. (2017). Pseudomonas baetica: pathogenicity for marine fish and development of protocols for rapid diagnosis. FEMS microbiology letters, 364(3), 10.1093/femsle/fnw286. 10.1093/femsle/fnw286

29. Chown S.L. et al. 2010. Adapting to climate change: a perspective from evolutionary physiology. Clim. Res. 43:3–15.

30. Hoffmann A.A. et al. 2003. Adaptation of Drosophila to temperature extremes: bringing together quantitative and molecular approaches. J. Therm. Biol. 28:175–216.

31. Cohen, J. M., Venesky, M. D., Sauer, E. L., Civitello, D. J., McMahon, T. A., Roznik, E. A., & Rohr, J. R. (2017). The thermal mismatch hypothesis explains host susceptibility to an emerging infectious disease. Ecology letters, 20(2), 184–193. 10.1111/ele.12720.

32. Cohen JM, Sauer EL, Santiago O, Spencer S, Rohr JR. 2020. Divergent impacts of warming weather on wildlife disease risk across climates. Science 370, abb1702. 10.1126/science.abb1702.

33. Hickey, D. A., & Singer, G. A. (2004). Genomic and proteomic adaptations to growth at high temperature. Genome biology, 5(10), 117. 10.1186/gb-2004-5-10-117

34. Pounds, J. A., Bustamante, M. R., Coloma, L. A., Consuegra, J. A., Fogden, M. P., Foster, P. N., La Marca, E., Masters, K. L., Merino-Viteri, A., Puschendorf, R., Ron, S. R., Sánchez-Azofeifa, G. A., Still, C. J., & Young, B. E. (2006). Widespread amphibian extinctions from epidemic disease driven by global warming. Nature, 439(7073), 161–167. 10.1038/nature04246

35. Boyd IL, PH Freer-Smith, CA Gilligan, HCJ Godfray, The consequence of tree pests and diseases for ecosystem services. Science 342, 1235773 (2013).

36. Patz, J., Campbell-Lendrum, D., Holloway, T. et al. Impact of regional climate change on human health. Nature 438, 310–317 (2005). 10.1038/nature04188.

37. Cressler, C., McLeod, D., Rozins, C., Van Den Hoogen, J., & Day, T. (2016). The adaptive evolution of virulence: A review of theoretical predictions and empirical tests. Parasitology, 143(7), 915–930. doi:10.1017/S003118201500092X.

38. Fisher, M., Henk, D., Briggs, C. et al. Emerging fungal threats to animal, plant and ecosystem health. Nature 484, 186–194 (2012). 10.1038/nature10947.

39. Rohatgi A. (2015). WebPlotDigitizer (Version 3.9) [Computer software]. Retrieved from http://arohatgi.info/WebPlotDigitizer.

40. Viechtbauer, W. (2010). Conducting Meta-Analyses in R with the metafor Package. Journal of Statistical Software, 36(3), 1–48. 10.18637/jss.v036.i03.

41. Anderson RM, May RM.. Coevolution of hosts and parasites. Parasitology 1982;85:411–26. Pandey, A., & Dawson, D. E. (2019). The shapes of virulence to come. Evolution, medicine, and public health, 2019 (1), 3. 10.1093/emph/eoy037.

42. Hairston, N. G. 1989. Ecological experiments: purpose, design and execution. Cambridge University Press, Cambridge, UK.

43. Porras, M.F., Agudelo-Cantero, G.A., Santiago-Martínez, M.G. et al. Fungal infections lead to shifts in thermal tolerance and voluntary exposure to extreme temperatures in both prey and predator insects. Sci Rep 11, 21710 (2021). 10.1038/s41598-021-00248-z

44. Greenspan, S. E., Bower, D. S., Roznik, E. A., Pike, D. A., Marantelli, G., Alford, R. A., Schwarzkopf, L., & Scheffers, B. R. (2017). Infection increases vulnerability to climate change via effects on host thermal tolerance. Scientific reports, 7(1), 9349. 10.1038/s41598-017-09950-3

45. Dayana Senthamarai, M., Rajan, M. R., & Bharathi, P. V. (2023). Current risks of microbial infections in fish and their prevention methods: A review. Microbial pathogenesis, 185, 106400. 10.1016/j.micpath.2023.106400

46. Longdon, B. et al. The causes and consequences of changes in virulence following pathogen host shifts. PLoS Pathog. 11, e1004728 (2015).

47. Arias, P.A., N. Bellouin, E. Coppola, R.G. Jones, G. Krinner, J. Marotzke, V. Naik, M.D. Palmer, G.-K. Plattner, J. Rogelj, M. Rojas, J. Sillmann, T. Storelvmo, P.W. Thorne, B. Trewin, K. Achuta Rao, B. Adhikary, R.P. Allan, K. Armour, G. Bala, R. Barimalala, S. Berger, J.G. Canadell, C. Cassou, A. Cherchi, W. Collins, W.D. Collins, S.L. Connors, S. Corti, F. Cruz, F.J. Dentener, C. Dereczynski, A. Di Luca, A. Diongue Niang, F.J. Doblas-Reyes, A. Dosio, H. Douville, F. Engelbrecht, V. Eyring, E. Fischer, P. Forster, B. Fox-Kemper, J.S. Fuglestvedt, J.C. Fyfe, N.P. Gillett, L. Goldfarb, I. Gorodetskaya, J.M. Gutierrez, R. Hamdi, E. Hawkins, H.T. Hewitt, P. Hope, A.S. Islam, C. Jones, D.S. Kaufman, R.E. Kopp, Y. Kosaka, J. Kossin, S. Krakovska, J.-Y. Lee, J. Li, T. Mauritsen, T.K. Maycock, M. Meinshausen, S.-K. Min, P.M.S. Monteiro, T. Ngo-Duc, F. Otto, I. Pinto, A. Pirani, K. Raghavan, R. Ranasinghe, A.C. Ruane, L. Ruiz, J.-B. Sallée, B.H. Samset, S. Sathyendranath, S.I. Seneviratne, A.A. Sörensson, S. Szopa, I. Takayabu, A.-M. Tréguier, B. van den Hurk, R. Vautard, K. von Schuckmann, S. Zaehle, X. Zhang, and K. Zickfeld, 2021: Technical Summary. In Climate Change 2021: The Physical Science Basis. Contribution of Working Group I to the Sixth Assessment Report of the Intergovernmental Panel on Climate Change [Masson-Delmotte, V., P. Zhai, A. Pirani, S.L. Connors, C. Péan, S. Berger, N. Caud, Y. Chen, L. Goldfarb, M.I. Gomis, M. Huang, K. Leitzell, E. Lonnoy, J.B.R. Matthews, T.K. Maycock, T. Waterfield, O. Yelekçi, R. Yu, and B. Zhou (eds.)]. Cambridge University Press, Cambridge, United Kingdom and New York, NY, USA, pp. 33−144, doi:10.1017/9781009157896.002.

48. Rangwala, I., Sinsky, E., Miller, J. R. (2013). Amplified warming projections for high altitude regions of the northern hemisphere mid-latitudes from CMIP5 models. Environmental Research Letters, 8(2), 024040. 10.1088/1748-9326/8/2/024040.

49. Bennett, J.M., Sunday, J., Calosi, P. et al. The evolution of critical thermal limits of life on Earth. Nat Commun 12, 1198 (2021). 10.1038/s41467-021-21263-8.

50. Alster, CJ, Weller, ZD, von Fischer, JC. 2018. A meta-analysis of temperature sensitivity as a microbial trait. Glob Change Biol. 24: 4211– 4224. 10.1111/gcb.14342.

51. Garcia-Solache MA, Casadevall A. 2010. Global warming will bring new fungal diseases for mammals. MBio. 1 (1):e00061–10. 10.1128/mBio.00061-10.

52. Bosch, J., Carrascal, L. M., Durán, L., Walker, S., & Fisher, M. C. (2007). Climate change and outbreaks of amphibian chytridiomycosis in a montane area of Central Spain; is there a link?. Proceedings. Biological sciences, 274(1607), 253–260. 10.1098/rspb.2006.3713.

53. Friedman DZP, IS. Schwartz. Emerging fungal infections–New patients, new patterns, and new pathogens. J Fungi, 5 (3) (2019), p. 67, 10.3390/jof5030067.

54. Gadre, A., Enbiale, W., Andersen, L. K., &#38; Coates, S. J. (2022). The effects of climate change on fungal diseases with cutaneous manifestations: A report from the International Society of Dermatology Climate Change Committee. The Journal of Climate Change and Health, 6, 100156. 10.1016/J.JOCLIM.2022.100156.

55. Cohen, JM, Civitello, DJ, Venesky, MD, McMahon, TA, Rohr, JR. An interaction between climate change and infectious disease drove widespread amphibian declines. Glob Change Biol. 2019; 25: 927– 937. 10.1111/gcb.14489.

56. Evans, S. S., Repasky, E. A., & Fisher, D. T. (2015). Fever and the thermal regulation of immunity: the immune system feels the heat. Nature reviews. Immunology, 15(6), 335–349. 10.1038/nri3843.

57. Dittmar, J., Janssen, H., Kuske, A., Kurtz, J. and Scharsack, J.P. (2014), Heat and immunity: an experimental heat wave alters immune functions in three-spined sticklebacks (Gasterosteus aculeatus). J Anim Ecol, 83: 744–757. 10.1111/1365-2656.12175.

58. Graham, A. L., Allen, J. E., Read, A. F. (2005). Evolutionary Causes and Consequences of Immunopathology. https://doi.org/10.1146/Annurev.Ecolsys.36.102003.152622, 36, 373–397. 10.1146/ANNUREV.ECOLSYS.36.102003.152622.

59. Cleaveland S. Diseases of humans and their domestic mammals: pathogen characteristics, host range and the risk of emergence. Philos. Trans. R. Soc. Lond. B Biol. Sci. 2001;356:991–999.

60. Woolhouse M.E.J., Gowtage-Sequeria S. Host range and emerging and reemerging pathogens. Emerg. Infect. Dis. 2005;11:1842–1847.

61. Taylor LH, Latham SM, Woolhouse MEJ (2001) Risk factors for human disease emergence. Philosophical Transactions of the Royal Society of London. Series B: Biological Sciences 356: 983– 989.

62. Gillooly, J. F., Brown, J. H., West, G. B., Savage, V. M. & Charnov, E. L. Effects of size and temperature on metabolic rate. Science 293, 2248–2251 (2001).

63. Schaum C.-E., Buckling A., Smirnoff N. and Yvon-Durocher G. 2022Evolution of thermal tolerance and phenotypic plasticity under rapid and slow temperature fluctuations. Proc. R. Soc. B. 2892022083420220834. 10.1098/rspb.2022.0834.

64. Garamszegi L. Z., Climate change increases the risk of malaria in birds. Glob. Change Biol. 17, 1751–1759 (2011). 10.1111/j.1365-2486.2010.02346.x.

65. Altizer, S., Ostfeld, R. S., Johnson, P. T., Kutz, S., & Harvell, C. D. (2013). Climate change and infectious diseases: from evidence to a predictive framework. Science (New York, N.Y.), 341(6145), 514–519. 10.1126/science.1239401.

66. Smith, K.F., Acevedo-Whitehouse, K. and Pedersen, A.B. (2009), The role of infectious diseases in biological conservation. Animal Conservation, 12: 1–12. 10.1111/j.1469-1795.2008.00228.x

67. Rohr JR, Cohen JM. Understanding how temperature shifts could impact infectious disease. PLoS Biol. 2020; 18(11):e3000938. 10.1371/journal.pbio.3000938

68. Harvell, C. D., Mitchell, C. E., Ward, J. R., Altizer, S., Dobson, A. P., Ostfeld, R. S., & Samuel, M. D. (2002). Climate warming and disease risks for terrestrial and marine biota. Science (New York, N.Y.), 296(5576), 2158–2162. 10.1126/science.1063699

69. IPCC, 2021: Summary for Policymakers. In: Climate Change 2021: The Physical Science Basis. Contribution of Working Group I to the Sixth Assessment Report of the Intergovernmental Panel on Climate Change [Masson-Delmotte, V., P. Zhai, A. Pirani, S.L. Connors, C. Péan, S. Berger, N. Caud, Y. Chen, L. Goldfarb, M.I. Gomis, M. Huang, K. Leitzell, E. Lonnoy, J.B.R. Matthews, T.K. Maycock, T. Waterfield, O. Yelekçi, R. Yu, and B. Zhou (eds.)]. Cambridge University Press, Cambridge, United Kingdom and New York, NY, USA, pp. 3−32, doi:10.1017/9781009157896.001.

70. Arias, P.A., N. Bellouin, E. Coppola, R.G. Jones, G. Krinner, J. Marotzke, V. Naik, M.D. Palmer, G.-K. Plattner, J. Rogelj, M. Rojas, J. Sillmann, T. Storelvmo, P.W. Thorne, B. Trewin, K. Achuta Rao, B. Adhikary, R.P. Allan, K. Armour, G. Bala, R. Barimalala, S. Berger, J.G. Canadell, C. Cassou, A. Cherchi, W. Collins, W.D. Collins, S.L. Connors, S. Corti, F. Cruz, F.J. Dentener, C. Dereczynski, A. Di Luca, A. Diongue Niang, F.J. Doblas-Reyes, A. Dosio, H. Douville, F. Engelbrecht, V. Eyring, E. Fischer, P. Forster, B. Fox-Kemper, J.S. Fuglestvedt, J.C. Fyfe, N.P. Gillett, L. Goldfarb, I. Gorodetskaya, J.M. Gutierrez, R. Hamdi, E. Hawkins, H.T. Hewitt, P. Hope, A.S. Islam, C. Jones, D.S. Kaufman, R.E. Kopp, Y. Kosaka, J. Kossin, S. Krakovska, J.-Y. Lee, J. Li, T. Mauritsen, T.K. Maycock, M. Meinshausen, S.-K. Min, P.M.S. Monteiro, T. Ngo-Duc, F. Otto, I. Pinto, A. Pirani, K. Raghavan, R. Ranasinghe, A.C. Ruane, L. Ruiz, J.-B. Sallée, B.H. Samset, S. Sathyendranath, S.I. Seneviratne, A.A. Sörensson, S. Szopa, I. Takayabu, A.-M. Tréguier, B. van den Hurk, R. Vautard, K. von Schuckmann, S. Zaehle, X. Zhang, and K. Zickfeld, 2021: Technical Summary. In Climate Change 2021: The Physical Science Basis. Contribution of Working Group I to the Sixth Assessment Report of the Intergovernmental Panel on Climate Change [Masson-Delmotte, V., P. Zhai, A. Pirani, S.L. Connors, C. Péan, S. Berger, N. Caud, Y. Chen, L. Goldfarb, M.I. Gomis, M. Huang, K. Leitzell, E. Lonnoy, J.B.R. Matthews, T.K. Maycock, T. Waterfield, O. Yelekçi, R. Yu, and B. Zhou (eds.)]. Cambridge University Press, Cambridge, United Kingdom and New York, NY, USA, pp. 33−144, doi:10.1017/9781009157896.002.

